# Time for What? Dissociating Explicit Timing Tasks Through Electrophysiological Signatures

**DOI:** 10.1101/2023.06.07.543718

**Authors:** Fernanda D. Bueno, Anna C. Nobre, André M. Cravo

## Abstract

Estimating durations between hundreds of milliseconds and seconds is essential for several daily tasks. Explicit timing tasks, which require participants to estimate durations to make a comparison (time for perception) or to reproduce them (time for action), are often used to investigate psychological and neural timing mechanisms. Recent studies have proposed that mechanisms may depend on specific task requirements. In this study, we conducted electroencephalogram (EEG) recordings on human participants as they estimated intervals in different task contexts to investigate the extent to which timing mechanisms depend on the nature of the task. We compared the neural processing of identical visual reference stimuli in two different tasks, in which stimulus durations were either perceptually compared or motorically reproduced in separate experimental blocks. Using multivariate pattern analyses, we could successfully decode the duration and the task of reference stimuli. We found evidence for both overlapping timing mechanisms across tasks as well as recruitment of task-dependent processes for comparing intervals for different purposes. Our findings suggest both core and specialised timing functions are recruited to support explicit timing tasks.

## Introduction

Interval timing, the ability to estimate durations in the hundreds of milliseconds to seconds, is essential for several daily tasks – such as perceiving speech and music, deciding when to switch driving lanes or performing a dance routine. Although this ability is commonly treated as a single skill, recent studies have proposed that how we process temporal durations may depend on how we use this information (Lewis and Miall, 2003; Coull and Nobre, 2008; Breska and Ivry, 2016; van Wassenhove, Herbst and Kononowicz, 2019).

One organising principle of timing tasks is how temporal information will be used and measured (Shalev, Nobre and van Ede, 2019; Coull and Nobre, 2008). A major distinction is between implicit and explicit timing tasks (Coull and Nobre, 2008). Implicit timing tasks do not require participants to report temporal durations; however, temporal durations nevertheless impact performance on another task-relevant factor. For example, temporally predictive cues can facilitate sensory and motor performance (Rohenkohl, Cravo, Wyart and Nobre, 2012; Rohenkohl, Gould, Pessoa and Nobre, 2014, Nobre and van Ede, 2018). In explicit timing tasks, temporal durations are the main objective, and participants report them in some fashion. Convergent evidence from lesion and correlational studies suggests that different neural systems contribute to implicit vs. explicit timing functions (Coull and Nobre, 2008; Breska and Ivry, 2016). Dissociations have also been proposed to prevail within different types of implicit timing tasks, such that the neural basis of temporal expectations may depend on the source of the temporally predictive signals as well as the task demands (Nobre and Rohenkohl, 2014; Breska and Ivry, 2018; Shalev, Nobre and van Ede, 2018; Breska and Ivry, 2020; Breska and Ivry, 2021; Nobre and van Ede, 2023).

Although explicit timing tasks have traditionally been treated as a homogeneous category, how participants provide explicit reports can differ significantly across tasks. For example, in motor tasks, participants estimate durations to produce timed actions, whereas in perceptual timing tasks, participants estimate durations to evaluate or compare them with a reference. To determine whether explicit timing functions are dependent on shared neural systems or timing functions unique to specific systems, it is crucial to understand the degree to which neural patterns of duration estimation overlap across these different tasks.

Previous studies exploring the relationship between perceptual and motor temporal judgments have primarily relied on behavioural and fMRI measures. Recent meta-analyses of fMRI data (Wiener, Turkeltaub and Coslett, 2010; Nani et al., 2019; Naghibi et al., 2023) have found that while certain brain regions, such as the Supplementary Motor Area and insula, are commonly activated across tasks, there are also task-specific activation patterns. Behaviorally, a similar interaction of global and goal-directed processes is also found. Studies by Merchant and colleagues showed that behavioural performance correlated across motor and non-motor timing tasks while also finding that tasks that required motor responses led to differences compared to perceptual tasks (Merchant, Zarco, and Prado, 2008a; Merchant, Zarco, Bartolo and Prado, 2008b).

Although research using brain imaging and behavioural methods suggests a combination of general and goal-directed processes involved in perceptual and sensorimotor timing, each approach has limitations. While behavioural findings can offer insight into how we express our time estimations, they do not enable us to compare whether or how different temporal processing stages are affected by task requirements. Similarly, functional brain imaging can identify common and distinct brain areas activated depending on the task goals but without indicating whether the dynamical patterns of activations within and across regions are comparable.

Electrophysiological studies in humans using EEG can offer valuable additional insights into how intervals are encoded and how task demands can influence different temporal processing stages. To date, many studies have investigated event-related potentials and timing (for a review, see Kononowicz, van Rijn, and Meck, 2018), most focusing on a single task or comparing the activity between temporal and non-temporal tasks. Recent developments in multivariate pattern analysis (MVPA) have demonstrated that EEG data also contain rich spatial information that can be used to decode neural states (Stokes, Wolff and Spaak, 2015). These methods have started to be applied to temporal tasks, revealing how task goals and context can influence how time is encoded in brain activity (Bueno and Cravo, 2021; Damsma, Schlichting, and van Rijn, 2021).

In this study, we investigated whether multivariate analysis of time-resolved EEG signals can distinguish between explicit timing of durations in tasks stressing perception vs. action. We designed an experiment in which participants viewed a reference interval in each trial and, in different blocks, had to reproduce the duration or compare it to a probe interval. The visual stimulus and set of durations/intervals were identical between conditions, enabling us to explore how EEG signals during and after the reference interval were modulated by the stimulus duration, task, and their interaction. With this approach, we aimed to advance the understanding of the neural mechanisms underlying perceptual and motor timing and how they differ and overlap.

## Materials and Methods

### Data availability

All materials from this study (task, analysis code, and raw and pre-processed data) will be available upon paper publication.

### Participants

The experimental protocol was approved by The Research Ethics Committee of the Federal University of ABC (UFABC), where the study took place. Study implementation followed approved guidelines and regulations (CAEE: 03607118.4.0000.5594).

Thirty-three human volunteers participated in the study after giving informed consent. Data from a final sample of twenty-nine volunteers were fully analyzed (mean age 23 y.o., age range of 18-35; 14 females). Data from the four additional participants were excluded from the analysis. The reasons included: loss of data due to an energy blackout during the experiment; loss of data due to computer memory issues during data collection; computer memory issues and excessive noise during data collection; and excessive data loss (19%) during artefact removal (see below). All participants had normal or corrected vision and declared being free from psychological or neurological diseases.

### Stimuli and Procedures

The experiment consisted of two computerized tasks presented in different blocks, temporal discrimination and temporal reproduction (Fig. 1a), combined with EEG recordings. The stimuli were presented using Psychtoolbox (Brainard, 1997) v.3.0 package in Octave on a ViewPixx monitor with a vertical refresh rate of 120 Hz, placed at approximately 52 cm from the participant. Responses were collected via a response box with nine buttons (DirectIN High-Speed Button; Empirisoft). In the discrimination task, participants used the index fingers of both hands to respond with the extreme left and extreme right buttons. In the reproduction task, they used the right index finger and the extreme right button.

**Fig 1.**
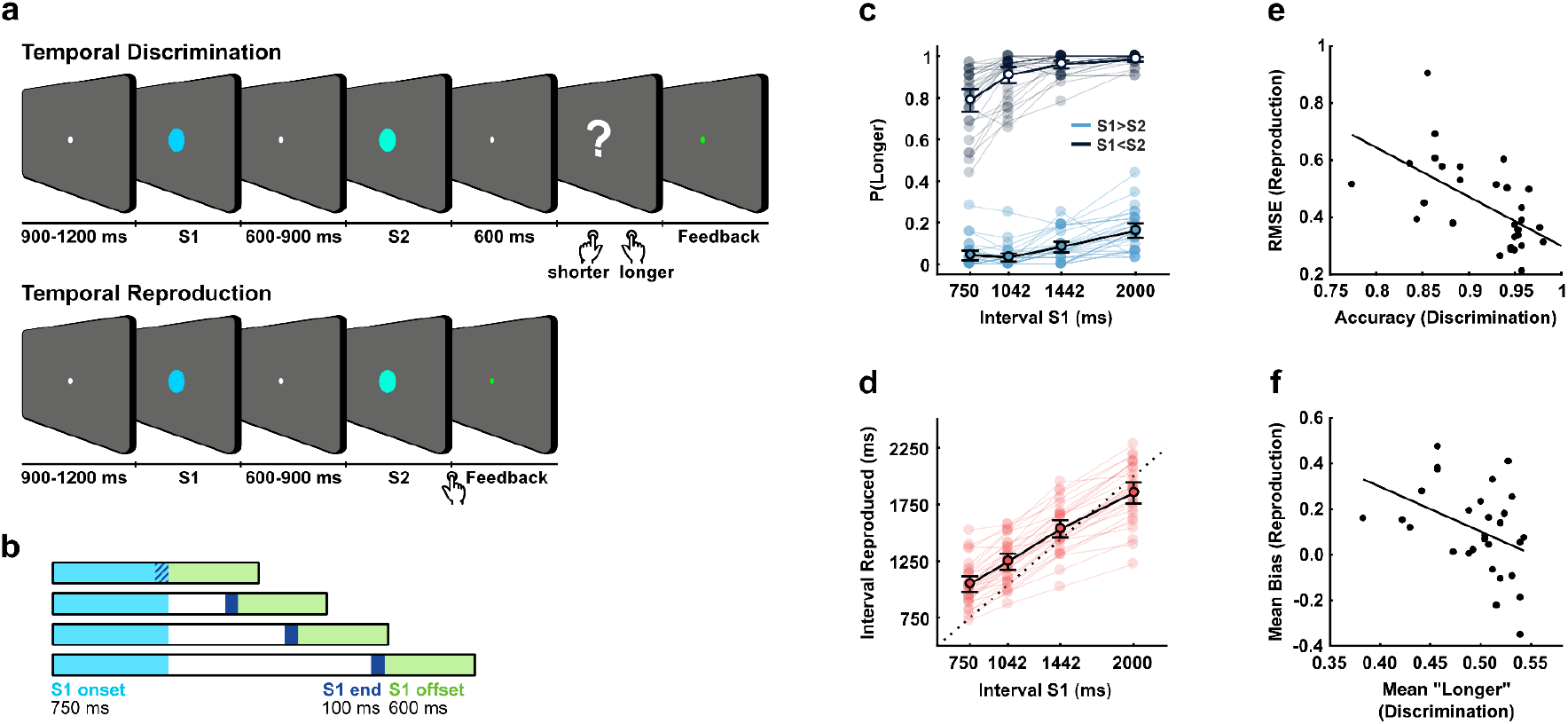
Experimental Design and Behavioural Results. (a) Schematic illustration of the discrimination and reproduction tasks. (b) Schematic illustration of the three segments used in the analysis, all relative to S1. The light blue area represents the period from 0 to 750 ms from S1 onset, whereas the dark blue area represents the final 100 ms before offset for all four sample intervals. The green area depicts the period analysed after S1 offset (c) Behavioural results for the discrimination task. The proportion of ‘longer than S1’ responses according to S1 interval (750, 1042, 1442, or 2000 ms) and whether S2 was shorter (light blue) or longer (grey) than S1. (d) Behavioural results for the reproduction task depict the mean of reproduced durations for each S1 interval. For (c) and (d), the light-colored circles depict each participant’s values, whereas black circles depict mean results for all participants. Error bars represent the standard error of the mean. (e) Correlation between the accuracy of responses (discrimination) and RMSE (reproduction) (robust skipped Spearman correlation, r = -0.59, 95% CI between -0.76 and -0.31; CI stands for Confidence interval). (f) Correlation between the mean of “longer” (discrimination) and mean bias (difference between reproduced and reference interval) (r =-0.40, 95% CI between-0.67 and -0.06). In (e) and (f), circles depict individual values and the line represents the correlation between values.

The experiment was divided into 16 short blocks of 32 trials each. Discrimination and reproduction tasks alternated from one block to another, and their order was counterbalanced between participants. A written cue presented for two seconds instructed and reminded participants about the task before each block started. The word ‘JULGAMENTO’ preceded the discrimination task (‘judgment’ in Portuguese), and ‘REPRODUÇÃO’ preceded the reproduction task (‘reproduction’ in Portuguese). For both tasks, trials consisted of two visual stimuli (filled circles of 1 visual-degree radius) presented sequentially on a grey screen.

Before each trial started, there was a blank screen (grey background) with a white fixation point at the centre (1/6 visual-degree radius) that could last between 900 and 1200 ms. The first stimulus (S1) was a light-blue-filled circle that could last one of four possible logarithmically spaced intervals: 750, 1042, 1442, or 2000 ms. The four intervals occurred equally frequently over the experiment. The order of the intervals presented at S1 was random, with the constraint that a given interval could not occur in more than three consecutive trials. After the first stimulus, another blank screen with a fixation point was shown (600 to 900 ms). We categorized this segment as the S1-offset epoch.

The second visual stimulus differed according to the task. In the temporal-discrimination blocks, the second stimulus (light-green-filled circle) lasted 40% less or 40% more than the first stimulus. After 600 ms from the offset of the second stimulus, a response screen appeared, prompting participants to indicate if the second stimulus was shorter (left button) or longer (right button) than S1. In the temporal-reproduction blocks, the second stimulus (S2) had the same light-green colour, but the participant controlled its duration. Participants were instructed to press the right button when they thought the reference (S1) duration had elapsed. The answer was considered correct if the reproduced duration was longer than half of the reference intervals and less than two times the reference. Feedback was provided for both tasks in every trial. The fixation point turned green if the answer was correct or red otherwise (for 350 ms duration).

Responses exceeding 11.2s (four times the longest interval for S2 in the discrimination task) were considered a timeout for both tasks. In such cases, no response was registered, negative feedback was provided, and the trial was considered lost. No trials were lost for any participant for reaching the timeout.

Before the experiment started, all participants completed a training session consisting of one block of each task. If the participant failed to achieve at least 75% accuracy in either task, the training session would be repeated up to three times. Only two participants required an extra training session.

### EEG recordings and pre-processing

EEG was recorded continuously from 64 ActiCap Electrodes at 1000 Hz by a QuickAmp amplifier (Brain Products). Data were high-pass filtered online (0.01 Hz) to avoid electrodermal fluctuations. All electrode sites were referenced to FCz and grounded to AFz. The electrodes were positioned according to the International 10-10 system, except for the TP9 and TP10 electrodes in the earlobes. Additional bipolar electrodes recorded the electrooculogram. Data pre-processing was carried out using the FieldTrip toolbox for MATLAB (Oostenveld, Fries, Maris and Schoffelen, 2011). Offline filters were applied to the continuous data with a bandpass of 0.1 Hz to 30 Hz (Butterworth filter, order 3), and all data were re-referenced to the average activity of the earlobe electrodes and downsampled to 250 Hz. Channels with missing data due to problems in acquisition or channels with excessive noise were interpolated using the FieldTrip channel-repair function. Data from most volunteers required no interpolation (14 participants) or up to two channels interpolated. Two participants had 3, and one participant had four channels interpolated.

Independent component analysis (ICA) was performed for eye-movement artefact detection and rejection. Eye-related components were identified with the help of SASICA, available for FieldTrip (Chaumon, Bishop and Busch, 2015), by visually inspecting topographies and time series from each component. We used 9-second segments from S1 onset to identify eye movement-related components to be rejected in later analysis relevant segments. Using these long segments for the ICA, we excluded periods between blocks from the continuous EEG data in which participants could move and interfere with the electoral signal and detection of eye-related components. From this point forward, we used data from 62 channels, excluding bipolar and reference electrodes on the earlobes (TP9 and TP10).

The analyses focused on the processing of the first stimulus. Critically, the first stimulus in both tasks was identical, and participants were not required to make any responses during this phase of the trial. We segmented the data relative to S1 onset (from 150 ms before to 2700 ms after) and S1 offset (from 150 ms before to 1 s after). Baseline correction used the periods from 150 ms before reference stimulus onset for S1-onset epochs. and 50 ms before to 50 ms after offset for S1-offset epochs. Baseline correction for S1-offset epochs was calculated around the 100 ms (-50 ms to 50 ms) from reference stimulus offset to remove any potential contribution from the classification related to the preceding final segment before offset. We evaluated interval and task decoding during three critical periods: from 0 to 750 ms after S1 onset (since this was the shortest possible S1 duration and therefore present for all trials); at the mean of the last 100 ms before S1 offset; and from 0 to 600 ms after S1 offset (since this was the shortest interval following S1) (Fig. 1b). Trials were rejected from the analysis if segments exceeded 150 μV or the amplitude range was greater than 250 μV in any of the 62 channels. The same segmentation for S1 and the final 100 ms was used for rejecting noisy trials. The percentage of rejected trials was 2.3% (range between 0% - 11.1%) for S1-onset segments (or the last 100 ms segments) and 0.7% for S1-offset segments (0%-5.9%).

### Multivariate pattern analysis

We investigated differences between tasks and intervals in the time-resolved EEG using a supervised learning classification approach. We used a Linear Discriminant Analysis (LDA) as implemented in the MVPA-Light toolbox for MatLab (Treder, 2020) combined with custom scripts. Decoding performance was estimated using accuracy when the classification consisted of two labels (i.e., decoding which task participants were performing). For ease of interpretation, classification accuracies were subtracted from chance levels (0.5 for two labels) for the two-label task classification. A multiclass classification was used for interval decoding: intervals were coded from 1 to 4, and performance was estimated using the mean absolute error (MAE). The MAEs were normalized using the following equation:

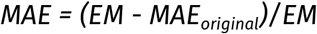

where EM is equal to 1.25 and stands for the expected mean of the absolute distances at the chance level (the sum of all absolute distances between the four labels, which is 20, divided by the number of all possible combinations of real and predicted levels, which is 16). After normalization, if the classification is perfect, the maximum result is one, and zero reflects chance-level performance. Similar results were obtained when classification accuracy was used.

For the time-resolved classification, data from individual trials were smoothed using a moving average with a 40-ms window. We used a four-fold cross-validation for task classification (reproduction or discrimination). Each fold consisted of eight blocks, four of each task. For example, in the first fold, the test data consisted of blocks 1 to 8 (four blocks of each task), and the training data consisted of the remaining blocks (9 to 32, twelve blocks of each task). In the second fold, test data consisted of blocks 9 to 16, and so on successively. Four-fold cross-validation was also used for interval classification within a task, each with two blocks of the task. For example, in the first fold, the test data consisted of the first four blocks of a given task, and the training data consisted of the other twelve blocks; in the second fold, the test data consisted of blocks 5 to 8 of that task, and so on successively. Lastly, we also performed a between-task classification for intervals, with training data consisting of 12 blocks of a task and the test data consisting of the other 4 blocks of the other task, following the same logic as the within-task classification.

At each time point, all 62 EEG sensors were used as features. The analysis was conducted at each time point (4 ms apart after downsampling). We evaluated the accuracy or MAE by permutation tests with the *tmax* statistic (Groppe, Urbach and Kutas, 2011), which corrects for multiple comparisons. All tests were one-sided t-tests compared with zero with an alpha level of 5% for significance level. For time point-by-point classification accuracy measures, we considered time points in which the t-values exceeded the empirical critical t calculated in the permutation test. Whenever relevant, we show univariate F-values for differences between experimental conditions to illustrate effects.

## Results

### Behavioral Results

Participants performed well on both tasks. Using the feedback levels as a measure of accuracy, the mean accuracy for the discrimination task was 91.6%, ranging between 77.3% and 98.0%. The mean accuracy for the reproduction task was 95.7%, ranging between 84.8% and 100%.

In the discrimination task (Fig. 1c), participants more often responded “longer” when S2 was longer than S1, as expected (F(1,28) = 1845.78, p < .001). Additionally, there was a higher proportion for “longer” responses for longer durations of S1 (F(1.741, 0.462) = 59.90, p < .001, Greenhouse-Geisser sphericity correction). Finally, we also found a significant interaction between these factors (F(2.091, 0.081) = 14.77, p < .001, Greenhouse-Geisser sphericity correction), indicating that the effect of duration on bias was stronger on both extremes intervals..

For the reproduction task (Fig. 1d), participants systematically produced longer responses as S1 duration increased. However, they tended to underestimate longer durations and overestimate shorter ones. The linear regression between the sample and reproduced intervals shows positive slopes but lower than one (95% Confidence Interval: 0.58 to 0.71, mean 0.65). Despite performing the task well, the pattern of participants’ responses followed the pervasive central tendency effect (Jazayeri and Shadlen, 2010).

We compared performance between tasks using two different approaches. In a first analysis, we investigated whether participants with a higher accuracy in discrimination also performed better in the reproduction task, as measured by the root-mean-square error (RMSE). The RMSE measures how far the interval reproductions are from the presented interval, with larger values indicating worse performance. Participants who performed better in one task were also better in the other task. We used a robust skipped correlation to protect against bivariate outliers (Pernet, Wilcox and Rousselet, 2013) and observed a significant negative correlation between discrimination accuracy and reproduction RMSE (robust skipped Spearman correlation, r = -0.59, 95% Confidence Interval: -0.76 -0.31, Fig. 1e). Using Spearman’s correlation yielded equivalent results (Spearman correlation, r = -0.61, 95% CI: -0.76 -0.35).

In a second analysis, we investigated whether participants were consistent in under/overestimating the duration of S1 between the two tasks. We correlated the average “longer” responses in the discrimination task and the bias in the reproduction task (calculated as the average difference between reproduced and reference intervals). There was a significant negative correlation between these measures, indicating that participants who reproduced intervals as longer tended to judge the second interval as being “shorter” in the discrimination task (robust skipped Spearman correlation, r = -0.40, 95% CI: -0.67 -0.06, Fig. 1f). Using Spearman’s correlation yielded equivalent results (Spearman correlation, r = -0.40, 95% CI: -0.66 -0.01).

### Electrophysiological Results

#### Timing task modulates EEG signals during S1

The first analysis focused on the differences in the EEG signal across tasks. We analyzed the first 750 ms from S1 onset from all non-rejected trials. Figure 2a shows that the accuracy for classifying the task was statistically above chance level, with effects starting during the early stages of the reference intervals (critical t = 3.527). For illustrative purposes, we performed a mass univariate ANOVA^1^ with the factors Task (discrimination and reproduction) and Interval (four possible reference intervals) for each time point and channel (Fig. S1b in Supplementary Material). The topographies for the univariate F-values across sensors in Fig. 2b show that differences in the time-resolved EEG between tasks were captured by more frontal and central sensors, as shown in Fig. 2c. We did not expect any modulation according to the interval during this initial period of S1 (since differential intervals had not yet elapsed), so this analysis also served as a sanity check and there no significant effects of interval durations.

**Fig 2.**
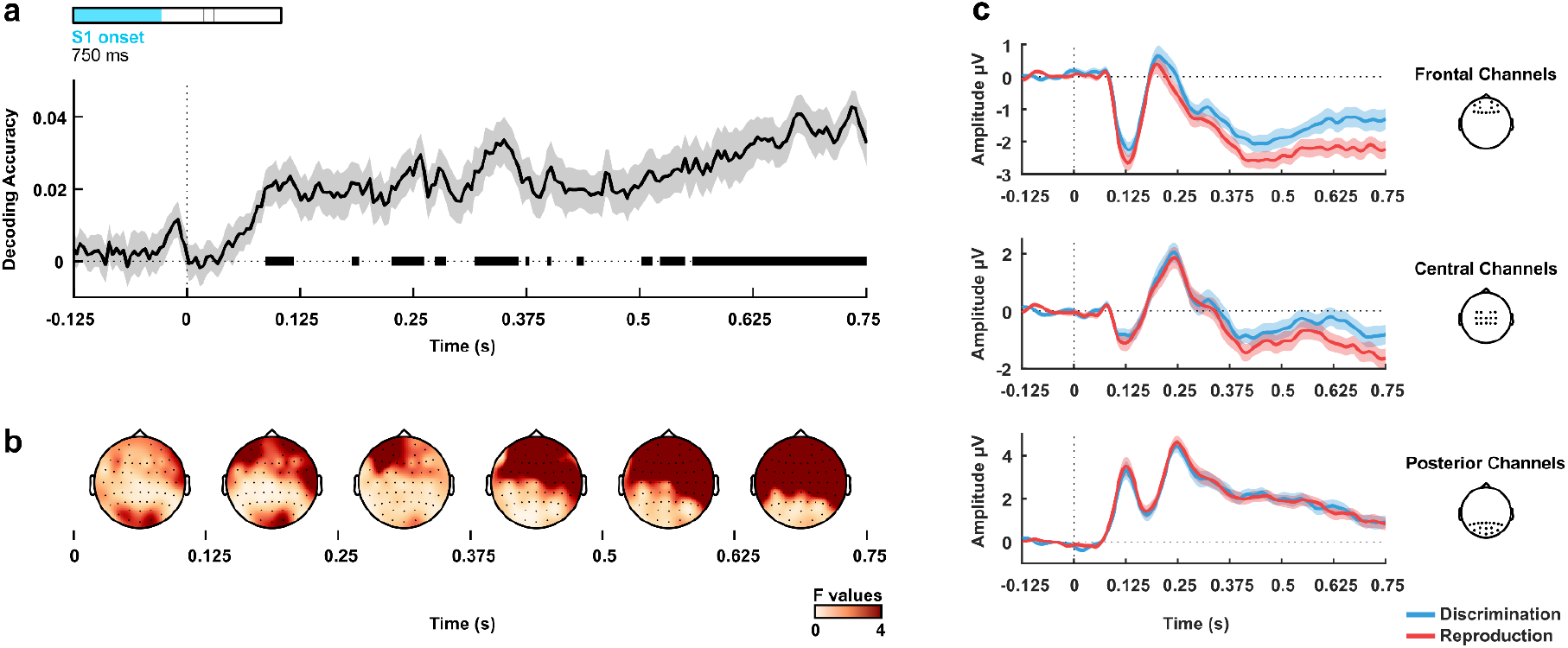
Task decoding during S1. (a) Task decoding accuracy above chance level (zero is chance level) throughout the first 750 ms from S1 onset (all valid trials). Bold horizontal lines indicate values significantly greater than zero (p<.05) from the permutation analysis. (b) Univariate F-values in 125 ms steps from 0 ms to 750 ms from stimulus onset. Differences across tasks were accentuated in frontal sensors at late periods. (c) Grand averages of the signal for the different tasks at frontal (Fp1, AF7, Fp2, AF8, F5, F3, F1, Fz, F2, F4, F6, AF4, AF3), central (FC3, FC1, FC2, FC4, C4, C2, Cz, C1, C3, CP3, CP1, CPz, CP2, CP4), and posterior channels (P8, P6, P4, P2, Pz, P1, P3, P5, P7, PO10, PO8, PO4, POz, PO3, PO7, PO9, O1, Oz, O2). The shaded areas represent the standard error of the mean.

#### Task and interval modulate activity at the end of the S1

Information about the duration of the interval is available at the end reference interval. We performed two separate analyses to test whether it was still possible to decode the task and, additionally, decode the duration of the reference interval. Both analyses used the average signal during the last 100 ms before S1 offset for each electrode as the dependent variable.

Classifying task yielded a significant above-chance decoding accuracy (t(28) = 6.887, p < .0001, Fig. 3a). Differences associated with the task were stronger in frontal and central sensors, as shown by the univariate F-values in Figures 3b and 3c.

**Fig 3.**
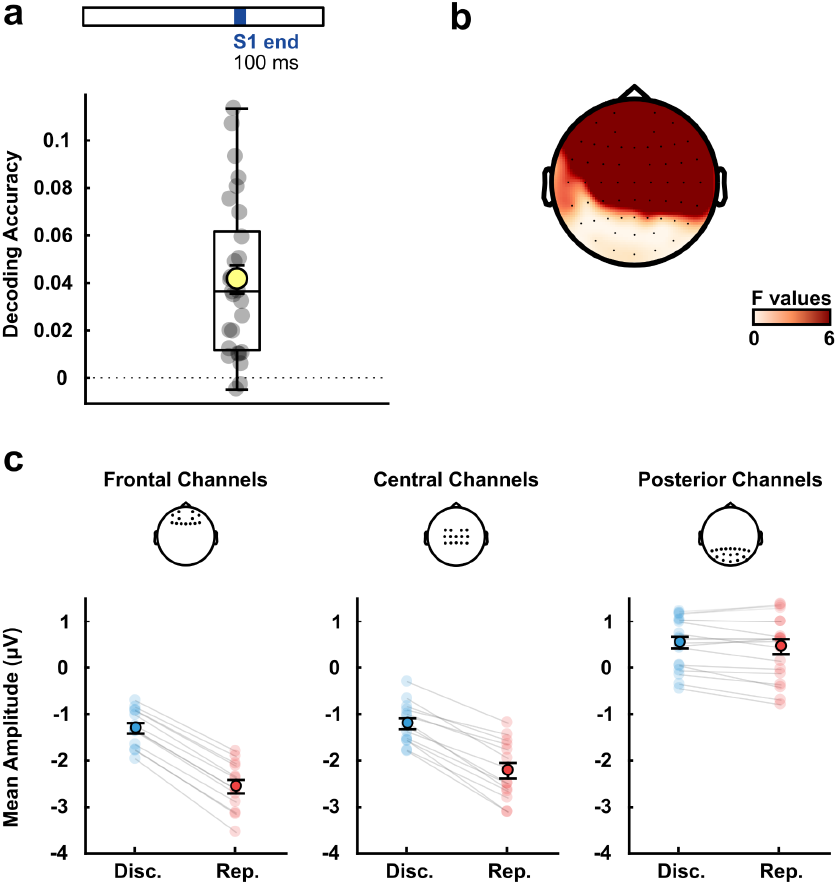
Task decoding in the last 100 ms of the reference stimulus. (a) Decoding accuracy above chance level (zero is chance level). The yellow circle depicts the mean accuracy for all participants and the smaller grey circles individual accuracies (b) Univariate F-values for task differences during the period. (c) Mean amplitude of the signal for the different tasks at frontal, central, and posterior channels (using the same channels as in Fig. 2). The light-coloured circles depict each participant’s values, whereas dark-coloured circles depict mean results for all subjects. Error bars represent the standard error of the mean.

We used both within- and between-tasks classification approaches to look at interval classification. In the within-task classification, training and testing data came from the same task (i.e., discrimination or reproduction). In the between-task classification, training used data from one task, and testing used data from the other. Significant MAE values resulted in all cases (Fig. 4a). Decoding was significant within tasks for discrimination (t(28) = 3.943, p<.001) and reproduction (t(28) = 4.042, p < 0.001). Between tasks, decoding was significant when training on reproduction and testing on discrimination (t(28) = 4.939, p < .0001) as well as when training on discrimination and testing on reproduction (t(28) = 6.602, p < .00001). In contrast to classifying the task, information about durations was most pronounced in central-parietal sensors (Fig. 4b and c).

**Fig 4.**
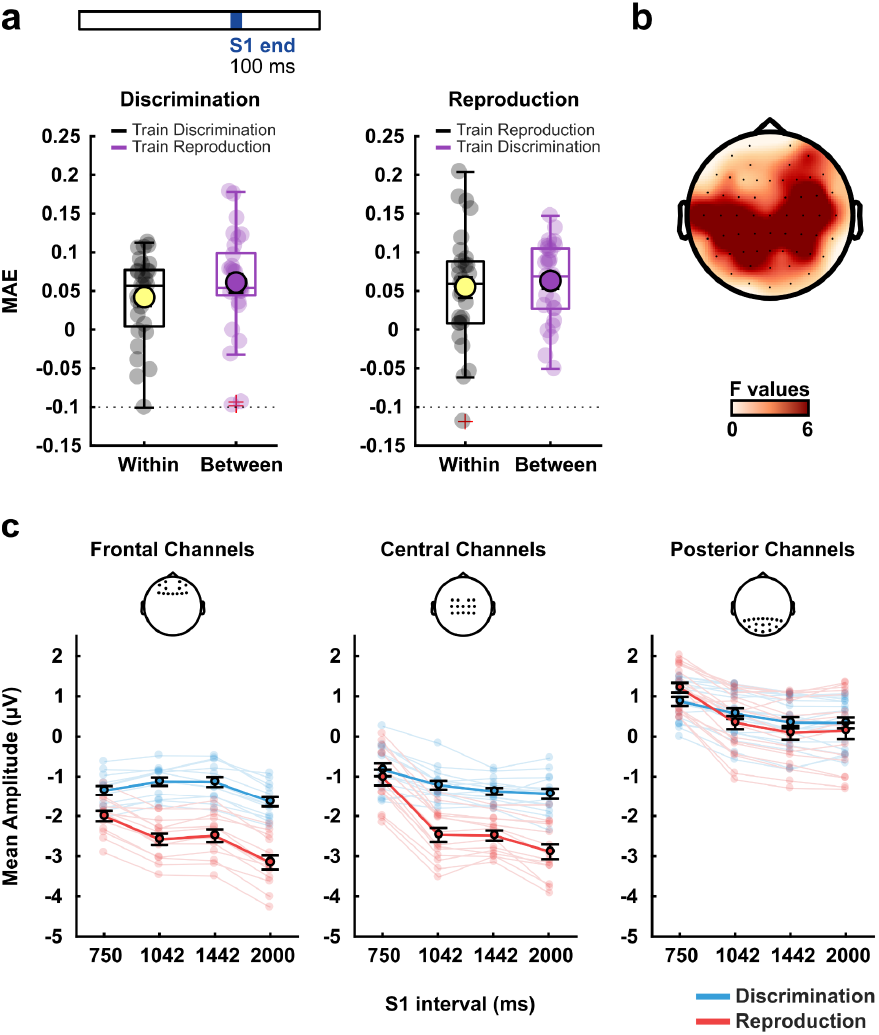
Time-interval decoding in the last 100 ms of the reference stimulus. (a) Decoding accuracy for the duration within and between discrimination and reproduction tasks. The yellow and dark purple circles depict the mean accuracies for all participants, and the grey or light purple smaller circles represent individual accuracies (b) Univariate F-values for interval differences during the period. (c) Mean signal amplitude for the different intervals and tasks at frontal, central, and posterior channels (using the same channels as in Figures 2 and 3). For (a) and (c), the light-coloured circles depict each participant’s values, whereas the dark-coloured circles depict mean results for all participants. Error bars represent the standard error of the mean.

#### Task and interval modulate activity after S1 offset

In a second analysis, we investigated if information about task and interval could also be found in the EEG signal after S1 offset. During this period, participants needed to maintain information about the reference interval to respond to the second stimuli. After S1 offset, we found that EEG activity is modulated by both task and duration.

The MVPA classifier decoded which task participants performed early after S1 offset (Fig. 5a, critical t = 3.467). Importantly, the data for this analysis were baseline corrected around the S1 offset, eliminating the effect reported for the last 100 ms of S1 task classification. Figures 5b and 5c show the evolution of EEG differences driven by task as illustrated by univariate F-values and the average signal at distinct electrodes.

**Fig 5.**
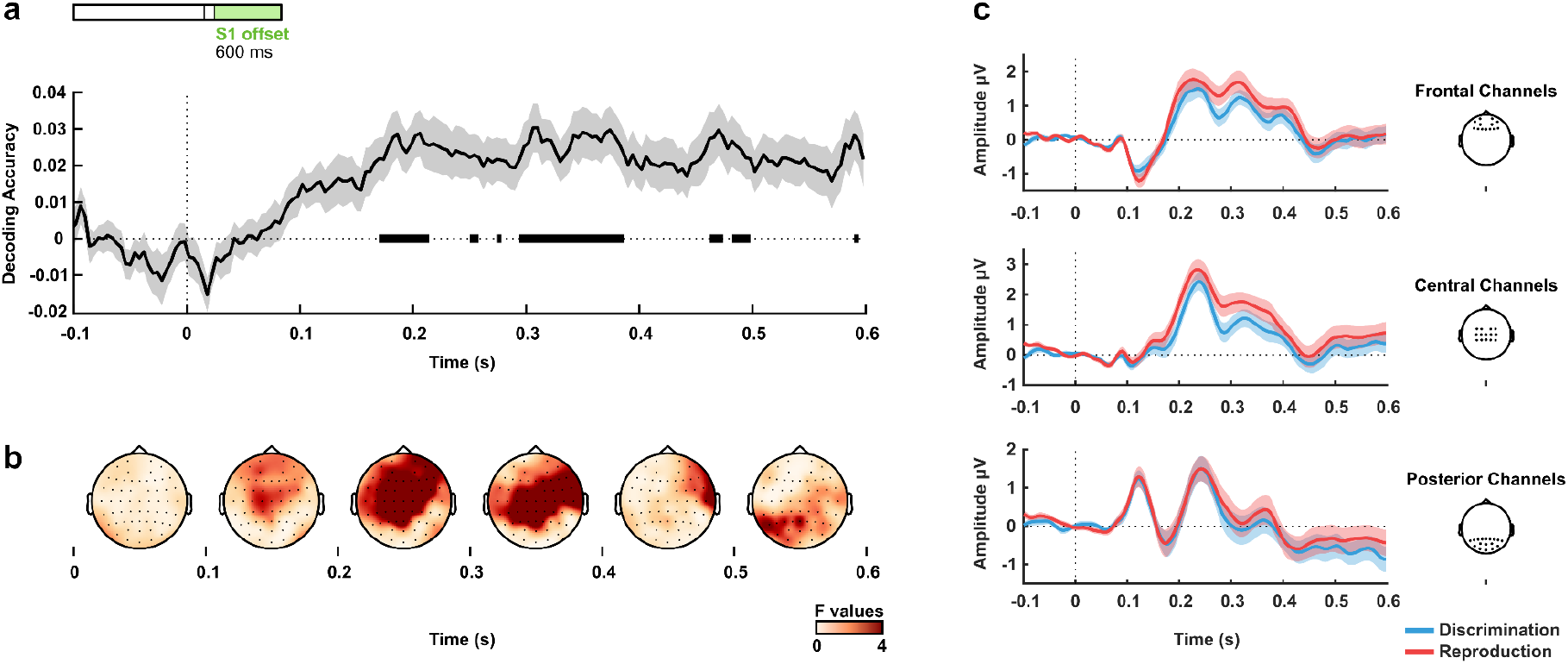
Task decoding after S1 offset. (a) Task decoding accuracy above chance level (zero is chance level) after S1 offset. Bold horizontal lines indicate values significantly greater than zero (p<.05) from the permutation analysis. (b) Univariate F-values in 125 ms windows from 0 to 600 ms from stimulus offset. Task-related differences were more pronounced at central electrodes. (c) Grand averages of the signal for the different tasks at frontal, central, and posterior electrodes. The shaded areas represent the standard error of the mean.

Decoding of intervals was done within and between tasks. Above-chance decoding levels were observed in both (Fig. 6). For the within-task classification, decoding was significant within tasks for discrimination (critical t = 3.563) and reproduction (critical t = 3.494). Between tasks, decoding was significant when training on reproduction and testing on discrimination (critical t = 3.578) and when training on discrimination and testing on reproduction (critical t = 3.577). Interestingly, looking at Figures 6b and 6c, two patterns emerge concerning the coding of intervals. An early posterior effect (100 to 300 ms) in which the amplitude is lower for shorter intervals and a later frontal effect (>300 ms) in which shorter intervals show a higher amplitude.

**Fig 6.**
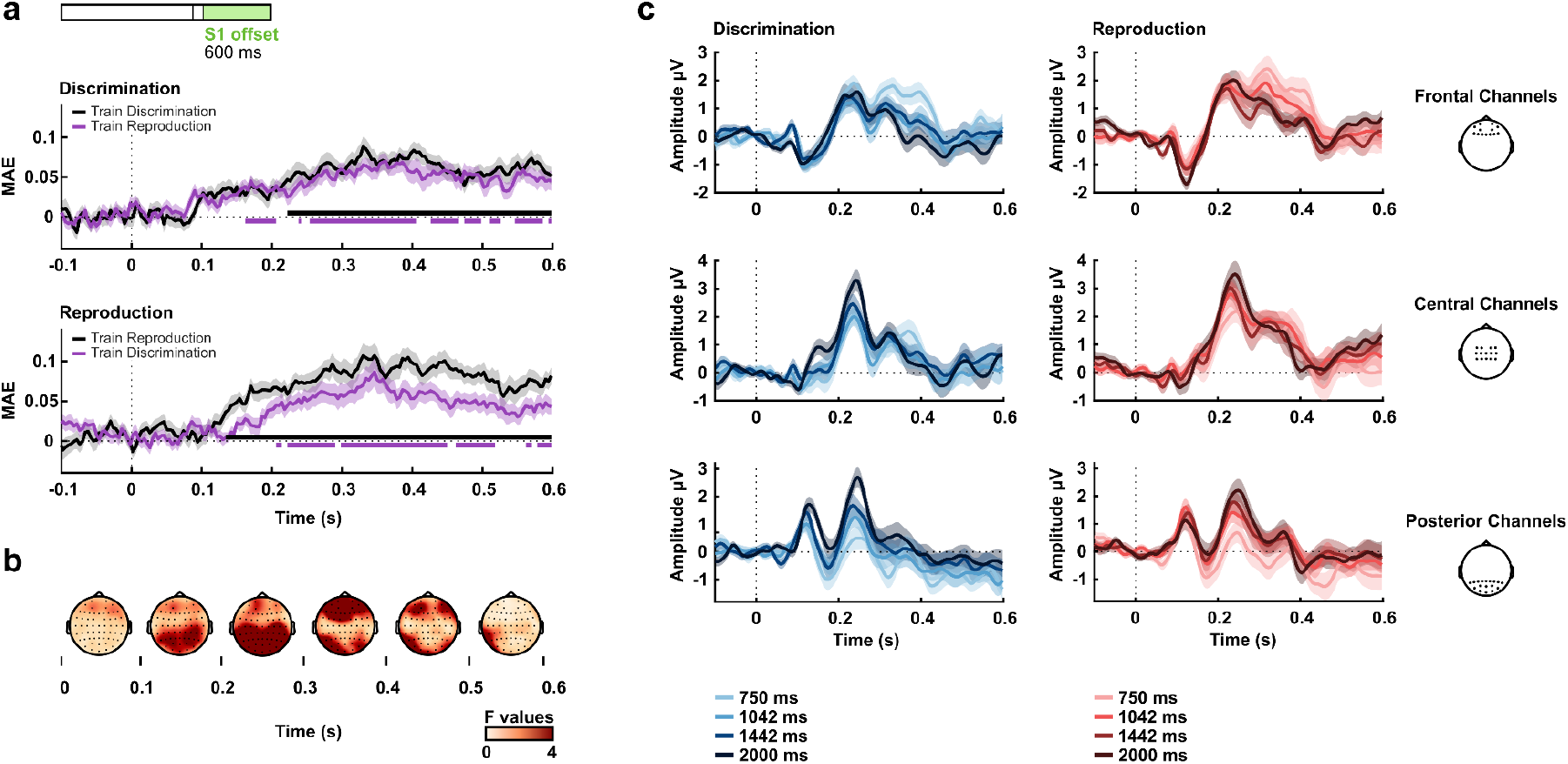
Time-interval decoding after S1 offset. (a) Decoding accuracy above chance level in the segment after S1 offset. Bold horizontal lines indicate values significantly greater than zero (p<.05) from the permutation analysis. (b) Average univariate F-values in 100 ms from 0 to 600 ms from stimulus offset. Differences related to duration were more pronounced at posterior electrodes at early periods (200 to 300 ms) and frontal at late periods (300 to 500 ms). (c) Grand signal averages for the different duration and tasks at frontal, central, and posterior electrodes (same channel separation as in Figures 2-5). The shaded areas represent the standard error of the mean.

#### Task-dependent time coding after S1 offset

Our final analysis investigated whether intervals are differently encoded across tasks. So far, we have shown that both task and time modulate EEG activity. Our previous analyses, in which time was trained and tested in different tasks, emphasised the commonalities for interval coding between tasks. In this last analysis, we investigated whether revealing selective, task-dependent processing of temporal intervals was also possible.

Our strategy was to use the difference in amplitude from the longest to the shortest interval and test whether MVPA can distinguish whether these differences come from performing different tasks. To increase the number of trials for the classifier, we created pseudo-trials comprising these differences. For each pseudo-trial, we draw four trials from the 2000-ms interval (longest) and four from the 750-ms interval (shortest) without replacement. We averaged the signal for each interval and the difference between the longest and shortest signal. Pseudo-trials were created using the same four-fold cross-validation strategy, so training and test pseudo-trials came from different folds. In the final step, a similar task classification was used on the pseudo-trials, investigating whether the classifier could successfully decode the task from the different subtractions. Creating pseudo-trials and classification was repeated 100 times for each participant, and the average accuracy was estimated.

Task-dependent differences in duration processing appeared after the offset of S1. No differences occurred during the last 100 ms of each interval (mean accuracy of 0.0076, standard error of the mean: 0.011; test difference from zero: t(28) = 0.707, p = 0.486). Differences after S1 offset are shown in Figure 7a. The periods in which the MVPA accuracy was above chance (critical t = 3.413) overlapped with periods in which we also observed common interval-duration coding between tasks. Figures 7b (discrimination) and 7c (reproduction) show the average differences from the longest to the shortest interval for discrimination and reproduction tasks, respectively, when the classifier accuracy was significantly above chance (from 378 ms to 414 ms). Figure 7d depicts the difference between discrimination and reproduction topographies (discrimination - reproduction).

**Fig 7.**
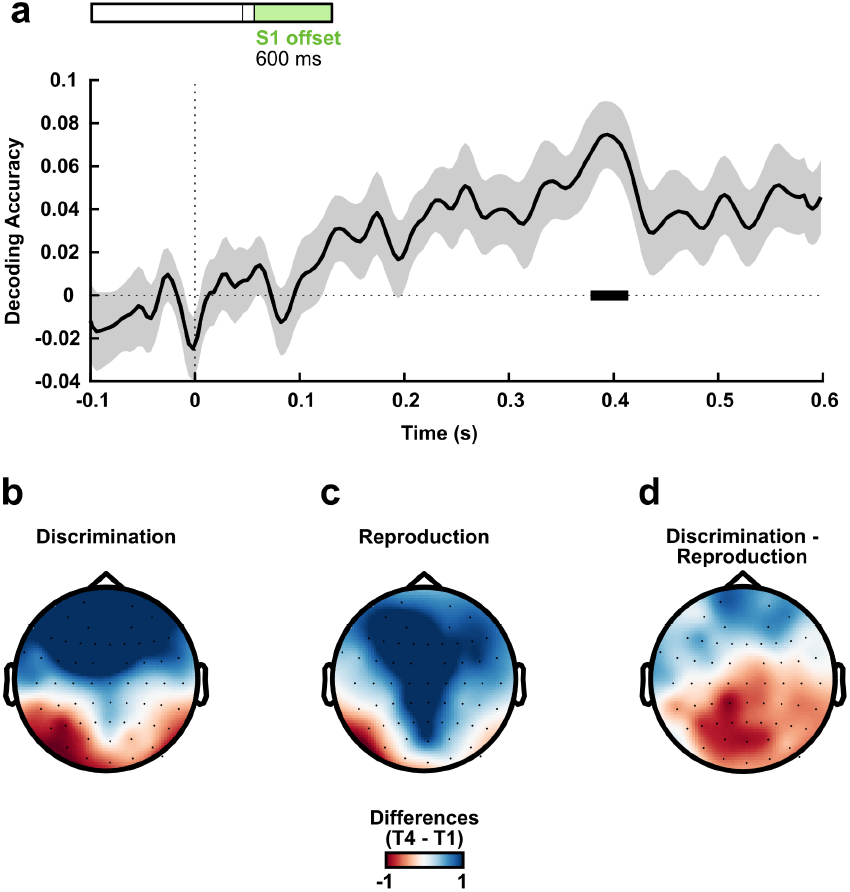
Task decoding after S1 offset on pseudo-trials isolating differences between shortest and longest references. (a) Decoding accuracy above chance level in the segment after S1 offset. Bold horizontal lines indicate values significantly greater than zero (p<.05) from the permutation analysis. (b) and (c) Mean differences from the longest to the shortest interval in discrimination and reproduction tasks, respectively, in moments where accuracy was significantly above chance (from 378 ms to 414 ms). (d) Mean difference between (b) and (c) topographies.

**Fig 8.**
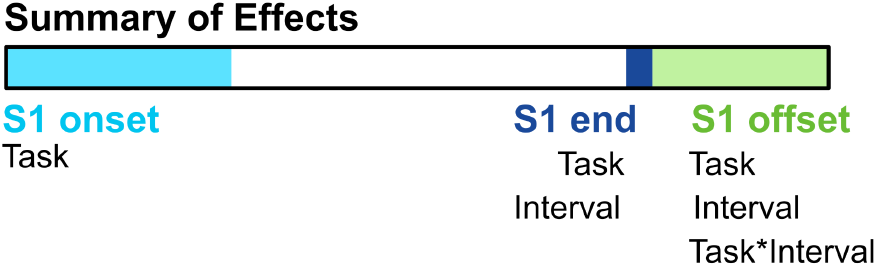
Summary of Effects. Schematic representation of the observed effects. We found task differences in brain activity on S1 onset, at the end of S1 and S1 offset. We found differences in brain activity across intervals at the end of S1 and S1 offset. Similar differences were both within and across tasks. Task-dependent differences in temporal intervals were found only after S1 offset.

## Discussion

Using multivariate pattern analysis, we revealed a set of overlapping neural signals for encoding the duration of temporal intervals for explicit perceptual judgment and motor reproduction complemented by distinct, task-selective processes for differentiating temporal durations. Despite analysing brain activity during periods with identical stimulation and timing demands, neural signals discriminating between the future task demands of perceptual judgment vs. motor reproduction were also prevalent across the various time scales analysed, ranging from early during the encoding interval to after the offset of the reference interval.

Task-specific EEG activity at the early stages of interval processing differed between encoding durations for subsequent perception- or action-based tasks, suggesting that timing unfolds in different neural contexts depending on future task demands. Within these different contexts, not only was it possible to decode timing within each task but decoding generalised between tasks, suggesting a common activity pattern that covaried with time. This common encoding of time was observed during both periods in which timing information was available - at the end of the reference interval and after its offset. After the interval offset, decoding of intervals was significant from early periods, including early differences over posterior sensors and later effects distributed over frontal sensors.

In a previous study (Bueno and Cravo, 2021), a similar pattern of duration encoding was found, but the effects were smaller and more spatially constrained, starting about 300 ms after the interval offset in frontal sensors. The similarity of the later effects between the two studies suggests there may have been a difference in data power or sensitivity. EEG correlates with duration was also reported after the offset of auditory-marked durations (Damsma, Schlichting, and van Rijn, 2021). In this case, effects started around 200 ms after the interval offset over central sensors. The different topographic distributions of the results suggest that the earliest effects after the offset of a reference stimulus may include sensory processing involving central sensors for the auditory task and posterior sensors for our visual task. Future studies could test for systematic variations in early timing-related signals depending on the sensory modality of the reference and test intervals.

Complementing our observation of robust cross-task decoding of temporal intervals, we also discovered that task demands impacted duration estimation. These task-dependent temporal encoding effects became apparent only after the offset of the reference interval. Unlike the common temporal encoding effects, the selective processes were not observed toward the end of the reference interval. The reason is unclear. Part of the explanation may rest on the need to derive the differences between stimulus durations between trials, which may have reduced the sensitivity and power of our analysis. Alternatively, the stimulus offset may be necessary to increase the readout of neural differences by acting as a perturbation that makes manifest differences in states that may otherwise remain encoded in “activity-silent” patterns of synaptic weights (Wolff, Ding, Myers and Stokes, 2015; Wolff, Jochim, Akyürek and Stokes, 2017). Moreover, after the interval offset, participants may start transforming and gating the encoded interval information to guide performance in the specific upcoming task. Such a process is likely to include the interaction between timing signals and proactive engagement of sensory vs. motor systems for perceptual judgment vs. motor reproduction, respectively. The differential encoding and maintenance of temporal intervals depending on how timing signals will be used are consistent with recent views of how working memory is flexibly coded depending on future use (Nobre and Stokes, 2019). Theoretical (Orhan and Ma, 2019) and empirical results in non-human primates (Warden and Mille, 2010) and fMRI in humans (Muhle-Karbe et al., 2017) have found that how information is maintained may depend on the expected use.

There has been an increasing interest in hybrid models for temporal processing, that propose a combination of local task-dependent areas interacting with partially distributed timing mechanisms (Merchant, Harrington, and Meck, 2013). Our findings align with the hybrid model proposal. We found a correlation in performance across tasks in our behavioural results and similar duration-dependent activity across tasks in our EEG results, suggesting common processing. However, we also found evidence for task-specific activity and an interaction of task-dependent activity and durations. These differences do not seem to be driven by tasks not being equally challenging, given that we found good performance in both and that we used short and intercalated task blocks to prevent learning or fatigue from modulating one task more than the other. However, we cannot completely rule out the possibility that general difficulty or motivation for the different tasks modulated EEG activity. On the other hand, the observed task-specific encoding of time shows how even these factors could modulate timing specifically.

In conclusion, our findings provide evidence that sensory versus motor demands may influence time encoding. These differences can have implications for other proposed categorisations of temporal processes, such as implicit versus explicit tasks. To gain a more comprehensive understanding of the neural mechanisms involved in temporal processing, future studies could employ higher spatial and spectral resolution techniques in humans and non-human animals to investigate these dimensions more systematically. Such research can yield valuable insights into the complex interplay of cognitive and neural factors that underlie our perception of time.

## Supplementary Material

**Fig S1.**
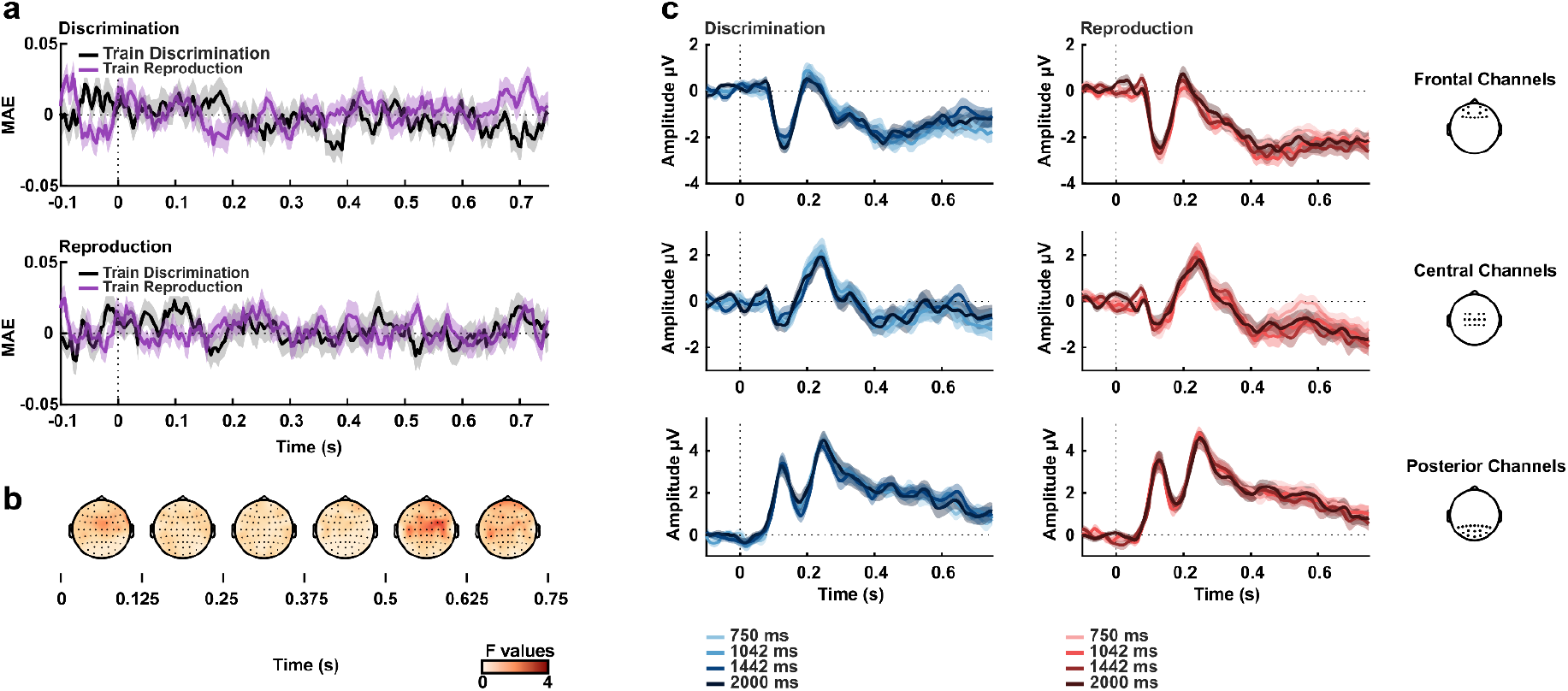
Time-interval during S1. (a) Decoding accuracy above chance level during S1 onset, throughout the first 750 ms from S1 onset (all valid trials) (b) Average univariate F-values in 125 ms from 0 to 750 ms from stimulus onset. (c) Grand signal averages for the different time intervals and tasks at frontal, central, and posterior electrodes (same channel separation as in Figures 2-6). The shaded areas represent the standard error of the mean.

## Footnotes

1 The ANOVA was performed for illustrative purposes. We plot F-values topographies for the factors Task and Interval in different figures.

